# The claustrum enhances neural variability by modulating the responsiveness of the prefrontal cortex

**DOI:** 10.1101/2024.09.27.615536

**Authors:** Huriye Atilgan, Ivan P. Lazarte, Adam M. Packer

## Abstract

The claustrum is recognized for its significant impact on various cognitive functions and its extensive connections with other brain regions, yet its functional role remains to be fully understood. Here, we utilized an optogenetic approach to investigate the claustrum’s influence on neuronal activity within the dorsal prefrontal cortex (dPFC) of mice. We conducted two-photon calcium imaging to assess dPFC cell responses during exposure to visual stimuli and widefield photostimulation of claustrum axons embedded in the dPFC. We identified three distinct subpopulations of neurons — sensory responsive, opto responsive, and opto-boosted cells — each exhibiting unique response dynamics to combined visual and optogenetic stimuli. Our findings reveal that stimulation of claustrum axons can normalize neuronal responsiveness, while enhancing neural variability, and significantly increasing network homogeneity. Training in a Pavlovian task showed that while enhanced variability with claustrum axon stimulation in neural responses persists, training does not further increase this variability but instead leads to greater network homogeneity. Additionally, we also performed claustrum axon silencing experiments that revealed that the claustrum may operate bidirectionally to maintain enhanced variability and homogeneity in the prefrontal cortex. These results highlight the crucial role of the claustrum in dynamically modulating dPFC activity, impacting both neuronal variability and network synchronization.

## Introduction

The claustrum and the prefrontal cortex (PFC) are tightly interconnected through a robust anatomical network (Torgerson et al., 2015; Wang et al., 2017; Zingg et al., 2018), hinting at a pivotal role for the claustrum in modulating executive functions governed by the PFC (Goll et al., 2015; Brown et al., 2017; Jackson et al., 2018; Liu et al., 2019; Chia et al., 2020; McBride et al., 2023; Atlan et al., 2024). While lesioning studies have not identified a single dysfunction associated with the claustrum, studies have unveiled intriguing insights into the claustrum’s involvement in a wide range of tasks (Atilgan et al., 2022; Jackson et al., 2020; Dillingham et al., 2017; Yin et al., 2016; Smythies et al., 2014). The claustrum, for example, has been linked to salience processing, highlighting its role in prioritizing stimuli and guiding attention allocation (Atlan et al., 2024; Chevée et al., 2022; Fodoulian et al., 2020; Smith et al., 2019; Atlan et al., 2018; Goll et al., 2015). Moreover, the claustrum is linked to integrating activity across brain regions during multimodal tasks, emphasizing its role in information processing and cognitive coordination (Shelton et al., 2024; Niu et al., 2022; Narikiyo et al., 2020; Krimmel et al., 2019; White & Mathur, 2018; Remedios et al., 2014; Crick & Koch, 2005; Hadjikhani & Roland, 1998, p. 199). Additionally, the claustrum is shown to activate during demanding tasks, supporting task switching and cognitive load management (Madden et al., 2022; Reus-García et al., 2021; Terem et al., 2020; White et al., 2020; Norimoto et al., 2020; Krimmel et al., 2019; Grasby & Talk, 2013). Despite these connections suggesting a significant role for the claustrum in cognitive function, the precise underlying mechanisms remain elusive.

Explorations of the claustrum’s impact on PFC cells have recently highlighted its inhibitory control, demonstrating a regulatory role in attention-based tasks and the complex mechanisms of claustrocortical communication (Jackson et al., 2018; Kim et al., 2016). This inhibitory influence reflects a dynamic interplay between the claustrum and cognitive processes. However, it’s important to recognize that the claustrum’s influence is not solely inhibitory. For instance, earlier research provides additional insights: Tsumoto and Suda (1982) reported bimodal excitatory responses in the visual cortex following dorsocaudal claustrum stimulation, with inhibition observed at longer latencies. Similarly, Cortimiglia et al. (1991) found that claustrum stimulation elicited low-latency excitatory effects on neurons in the medial oculomotor area of the cats, equivalent to the primate frontal eye fields, projecting to the superior colliculus. These findings underline the complex and multifaceted role of the claustrum in modulating PFC response to sensory inputs, highlighting both excitatory and inhibitory influences within neurocircuitry.

Building upon this foundation, our study aims to elucidate the claustrum’s contribution to PFC neural responsiveness, exploring its influence in both naive and trained states. By exploring the mechanisms governing claustrum-PFC interactions, we reveal how the claustrum influences sensory processing, enabling the brain to navigate and respond effectively to complex stimuli.

Our experimental approach involved recording from mice dPFC while modulating claustrum axons embedded within the PFC using two-photon microscopy. Mice were exposed to visual stimuli with and without photostimulation before and after visual stimulus-reward association training. Our findings suggest that the claustrum dynamically regulates PFC responsiveness to visual cues, acting as a modulator that can enhance or suppress sensory processing to optimize neural variability and network homogeneity. This modulation appears to be influenced by the baseline responsiveness in the PFC neurons and can be impacted by learning, highlighting the integral role of the claustrum in adjusting sensory processing and cognitive functions, enabling adaptive responses to internal and external demands.

## Results

### Claustrum axon photostimulation alters dPFC cell responsiveness

We employed an optogenetic experimental framework to elucidate how claustrum axons modulate responses in the dorsal prefrontal cortex (dPFC), (**Fig. 1A**). We monitored neural activity in dPFC in mice during exposure to visual stimuli, while photostimulating claustrum axons in the same area. Genetically modified mice expressed the calcium indicator GCaMP6s in CaMKII+ cells and the red-shifted opsin ChrimsonR (Klapoetke et al., 2014) in claustrum axons (**Fig. 1B-C**, see methods for our injection strategy). We employed two-photon microscopy in head-fixed mice for population calcium imaging. The mice were exposed to visual stimuli (“visual”), photostimulation (“opto”), and a combination of both visual stimuli and photostimulation (“visual+opto”) in various inter-trial intervals (**Fig. 1D**; 30 trials each, randomly presented). This setup facilitated the observation of neural dynamics at the single-cell resolution across a large field of view (580 x 580 µm) as we administered wide-field photostimulation (cranial window: ∼4000 µm diameter) to claustrum axons, thereby assessing the influence of claustrum axon photostimulation on dPFC cellular responses to visual stimuli.

**Figure 1:**
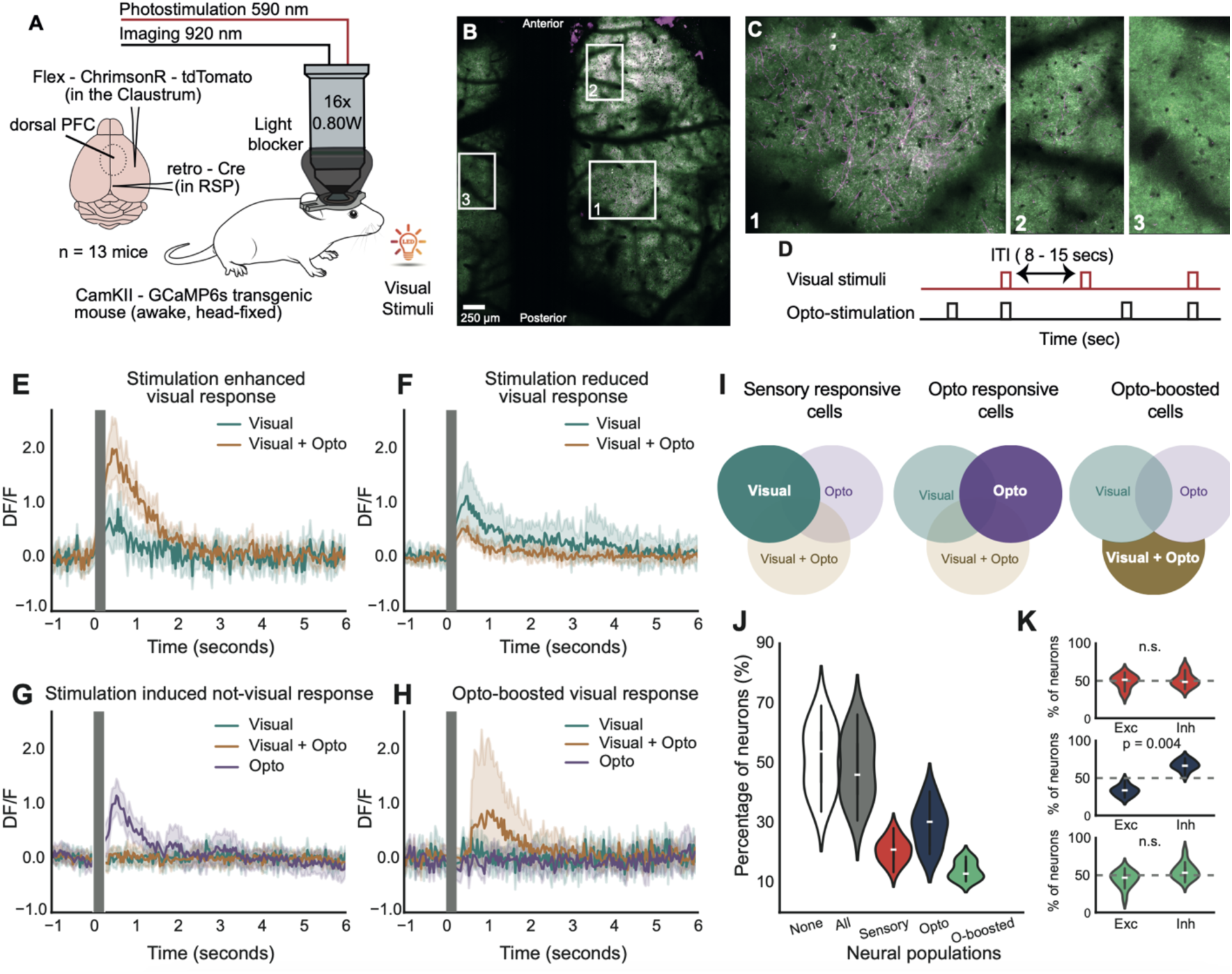
Claustrum axon photostimulation modulates dPFC cell responses **A)** Schematic of the experimental setup for two-photon imaging of dorsal PFC neurons expressing GCaMP6s in head-fixed, awake mice, with concurrent photostimulation of claustrum axons during visual cue presentation. **B)** Stitched two-photon image of the dorsal surface of the brain showing GCaMP-expressing neurons (green) and claustrum axons labelled with tdTomato (magenta) within the imaging field of view. **C)** Zoomed-in images from Panel B. Image 1 shows the embedded axons; image 2 shows a more anterior field of view; image 3 shows a contralateral field of view without any axon labelling as an outcome of the ipsi-hemisphere specific projections of the claustrum. **D)** Timing sequence of optogenetic stimulation and visual stimuli presentation across trials with variable inter-trial intervals (ITIs). **E-H)** Time course of neuronal calcium responses (DF/F) illustrating the modulatory effects of claustrum axon photostimulation on dPFC sample neuron responses to visual stimuli. **I)** Illustration of 3 subpopulations as sensory responsive cells population includes all cells responsive to visual cues, while opto responsive cells population includes all cells responsive to opto cues and opto-boosted cells population includes cells that do not respond to visual or opto stimuli presented in isolation, but only together. **J)** Violin plots for the distribution of dPFC neuron response across different subpopulations including all responsive (all) and none responsive (None) cells. **K)** Violin plots for the proportion of excitatory (Exc) and inhibitory (Inh) cells for each subpopulation. (n.s indicates not significant in Wilcoxon test)

Calcium responses revealed diverse changes in dPFC neurons following claustrum axon photostimulation. Some neurons exhibited enhanced responses to visual+opto stimuli (**Fig. 1E**), while others showed reduced responses under the same conditions (**Fig. 1F**). Notably, these effects were observed in both excited and inhibited responses (**Supp. Fig 1.2 A, B**), highlighting the high diversity of the claustrum axons’ impact on dPFC neuronal activity to visual stimuli.

A subset of neurons exhibited activity exclusively in response to opto stimuli, showing no discernible reaction to visual stimuli alone or when presented in combination with photostimulation (**Fig. 1G**). On the other hand, a distinct subset of cells demonstrated an interesting phenomenon where photostimulation effectively enhanced their responsiveness to visual stimuli, even though these cells did not exhibit any response when presented with visual stimuli or photostimulation alone (**Fig. 1H, Supp Fig. 1C**). These findings shows the complex and precise modulation of neuronal responses in the dPFC by claustrum axon photostimulation, highlighting its role in selectively enhancing or suppressing neural activity under varying condition.

Upon analyzing all the recorded cells, we found that 49% of the total recorded cells exhibited responsiveness (**Fig. 1J**, n = 24,575 from 49,212 recorded cells, n = 13 mice, 270 field of view), with responsiveness defined based on an independent t-test conducted 1 second before and after either visual, opto, or visual+opto stimuli trials (false discovery rate used for multiple correction, ⍺ = 0.05). Within this responsive cohort, 21% showed responsiveness to visual stimuli (n = 10,531) and were classified as *sensory responsive cells*, with 42% of these cells being exclusively responsive to visual stimuli. 29% responded to opto stimuli (n = 11,365) and were classified as *opto responsive cells*, among which 44% were exclusively responsive to opto stimuli. 3% of these cells (n = 1802) exhibited responsiveness to either visual or opto stimuli alone but not to visual+opto stimuli. Furthermore, 14% of the cells (n = 5783), were classified as *opto-boosted cells*, and displayed responsiveness to visual stimuli only when preceded by photostimulation, indicating a possible interaction between the visual stimuli and photostimulation that is not evident when each stimulus is presented alone.

To better understand how these combined subpopulations within the same network process visual and opto stimuli, we examined the balance between excitatory and inhibitory responses, which is crucial for neural network information processing (Froemke, 2015; Van Vreeswijk & Sompolinsky, 1996). This analysis revealed that opto-responsive cells uniquely exhibited a higher prevalence of inhibited responses, with an excitatory to inhibitory (E: I) ratio of 33:67 (**Fig. 1K**). When considering exclusively opto stimuli responsive cells within this subpopulation (excliding any cells responsive to visual or visual+opto stimuli), the E: I ratio becomes even more distinct, with the ratio changing from 33:67 to an even more pronounced 28:72. Notably, this trend was not observed in sensory-responsive and opto-boosted cells. The higher prevalence of inhibited responses observed in opto-responsive cells aligns with existing literature (Jackson et al., 2018) and suggests that optogenetic stimulation may indeed exert an inhibitory effect within the neural circuitry, particularly for cells not exhibiting any functional responsiveness. However, the absence of this trend in other subpopulations may indicate that the inhibitory effects of claustrum axon stimulation are more subtle or are potentially overridden by the stronger excitatory influence of visual stimuli, suggesting that inhibition is just one component of the complex and dynamic interactions within the dPFC neural circuitry.

### The claustrum modulates dPFC neuronal responses to enhance neural flexibility

To clarify the role of three identified subpopulations in the dPFC during claustrum axon photostimulation, we performed targeted analyses focusing on their individual response patterns to better understand their functional dynamics in sensory processing.

Initially, we focused on sensory responsive cells and, consistent with the literature, observed varying visual cues responses among dPFC cells (Wool et al., 2023; Barthas & Kwan, 2017; Moore & Armstrong, 2003), displaying different levels of excitation and inhibition (**Fig. 2A**). Upon visual+opto stimulation (**Fig. 2B**), these cell responses notably changed (**Fig.2C**), with excited responses in cells that were originally inhibited by visual stimuli alone and inhibited responses in those that were excited. To quantify this effect, we calculated the mean ΔF/F for 1500 ms after stimulation. The resulting histogram for visual+opto stimuli showed a significantly different distribution, where photostimulation shifted the bimodal distribution toward a Gaussian-like curve, indicating a normalising effect on neuronal responsiveness (**Fig. 2D**). At the population level, the absolute magnitude of neuronal responses to visual stimuli significantly decreased in the presence of photostimulation (**Fig. 2E**, *p* < 0.001, Kolmogorov-Smirnov test). To better understand how the distribution of neuronal responses was affected, we performed an empirical cumulative distribution function analysis, which revealed that a greater number of neurons were responsive when visual+opto stimuli were presented compared to visual stimuli alone (**Supplementary Fig. 2.1A**). This pattern suggests that, although the amplitude of individual responses decreased, the optogenetic intervention recruited a larger population of neurons, thereby enhancing distributed sensory processing with a balanced level of responsiveness.

**Figure 2:**
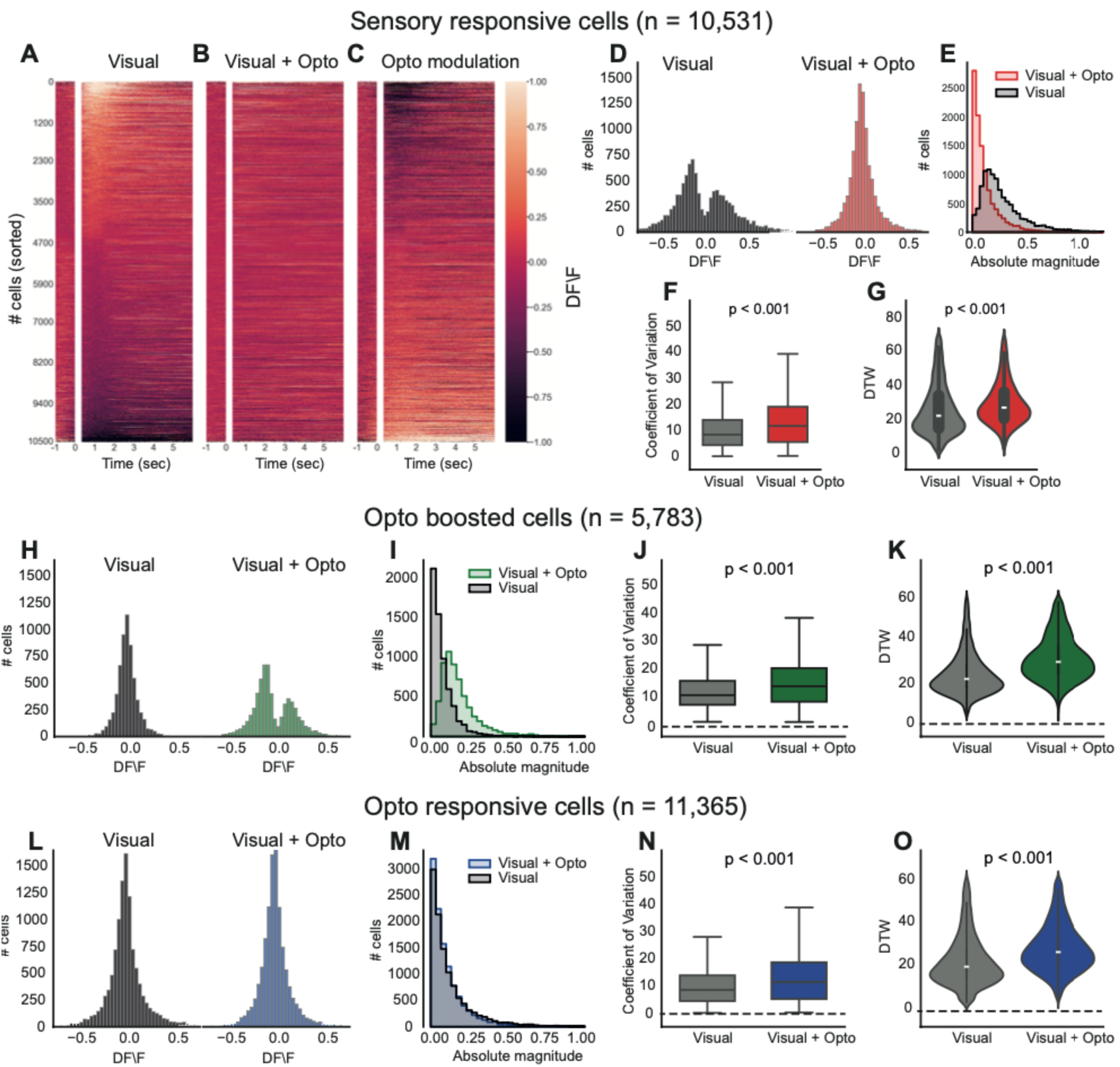
Claustrum axon modulation enhances neural flexibility in the dPFC A-C) Heatmap of neuronal activity over time for neurons under three different conditions: Visual (A), Visual + Opto (B) and their differences (C, opto modulation) **D)** Distribution of the DF/F for neurons under the conditions of visual and visual + opto for sensory responsive cells. **E)** Histograms representing the absolute magnitude of neuronal responses to visual and visual + opto stimuli for sensory responsive cells. **F)** Box plots comparing the coefficient of variation for neurons under visual and visual + opto conditions for sensory responsive cells**. G)** Violin plots illustrating the distribution of dynamic time warping (DTW) values under visual-only and visual + opto conditions. **H-K)** Similar to D-G, but for opto-boosted cells. **L-O)** Similar to D-G, but for opto responsive cells.

To explore the mechanisms underlying this distributed processing, we hypothesized that the claustrum might increase neural flexibility in the dPFC by elevating the variability in neuronal responses across repeated trials. We initially quantified this variability by examining the coefficient of variation, which showed a significant increase under visual+opto stimulation (**Fig. 2F**, *p* < 0.001, paired t-test). We then applied dynamic time warping (DTW) analysis—a technique that measures the similarity between two temporal signals—across trials for both visual and visual+opto stimuli. A higher DTW value indicates a greater disparity between two sequences, necessitating more adjustments to achieve alignment. Significantly larger DTW values for the visual+opto condition (**Fig. 2G**, *p* < 0.001, paired t-test) demonstrated increased variability when visual cues were combined with optostimulation, suggesting that the neurons are adapting their firing patterns more distinctly in each trial. This finding supports our hypothesis that claustral modulation enhances neural flexibility by increasing the variability of neuronal responses within the dPFC.

Subsequent analysis focused on the opto-boosted subpopulation, a unique subpopulation that did not respond to visual or optogenetic stimulation alone but showed activity when these stimuli were combined. The response distribution of these neurons shifted from a non-responsive state to a bimodal pattern of activation upon visual-optogenetic stimulation, as illustrated by histograms (**Fig. 2H-K**). Notably, there was a significant increase in the absolute magnitude of responses to visual+opto stimuli compared to visual stimuli alone (**Fig. 2I**, *p* < 0.001, Kolmogorov-Smirnov test). This suggests that photostimulation can unveil latent responsiveness in certain dPFC neurons, by making the neurons more excitable. This subpopulation also exhibited a higher coefficient of variation under visual+opto stimuli (**Fig. 2J**, *p* < 0.001, paired t-test) and more pronounced temporal variability in dynamic time warping analyses (**Fig. 2K**, *p* < 0.001, paired t-test) highlighting an enhanced flexibility in their response patterns when visual stimuli presented with optostimulation.

Although opto-responsive neurons do not show changes in the absolute magnitude of their response to visual cues (**Fig. 2M**, *p* = 0.192, paired t-test), they also exhibited increased variance in their neural response for visual+opto stimuli (**Fig. 2N**, *p* < 0.001, paired t-test) and DTW analysis (**Fig. 2O**, *p* < 0.001, paired t-test).

Despite the differing results observed in the first two subpopulations — where sensory-responsive cells exhibit normalized responses and opto-boosted cells reveal latent responsiveness—these outcomes collectively suggest that claustrum stimulation in the dPFC enhances the flexibility of neural responses. The observed increase in the coefficient of variation and pronounced temporal variability indicates greater variability in neuronal responses across all subpopulations when subjected to combined visual and optogenetic stimuli. Collectively, these findings suggest that the claustrum plays a crucial role in facilitating flexible and distributed processing mechanisms within the dPFC.

### Greater dPFC network homogeneity with claustrum axon activation

The greater response variation we observed in these subpopulations was consistent across all animals as well as in non-responsive cells (**Fig. 3A-C**), suggesting that optogenetic stimulation broadly affects the neural network beyond just the most noticeable responsive cells. This implies a widespread influence of the claustrum on the dPFC, potentially affecting even those neurons that do not typically react strongly to visual and/or optogenetic stimuli alone.

**Figure 3:**
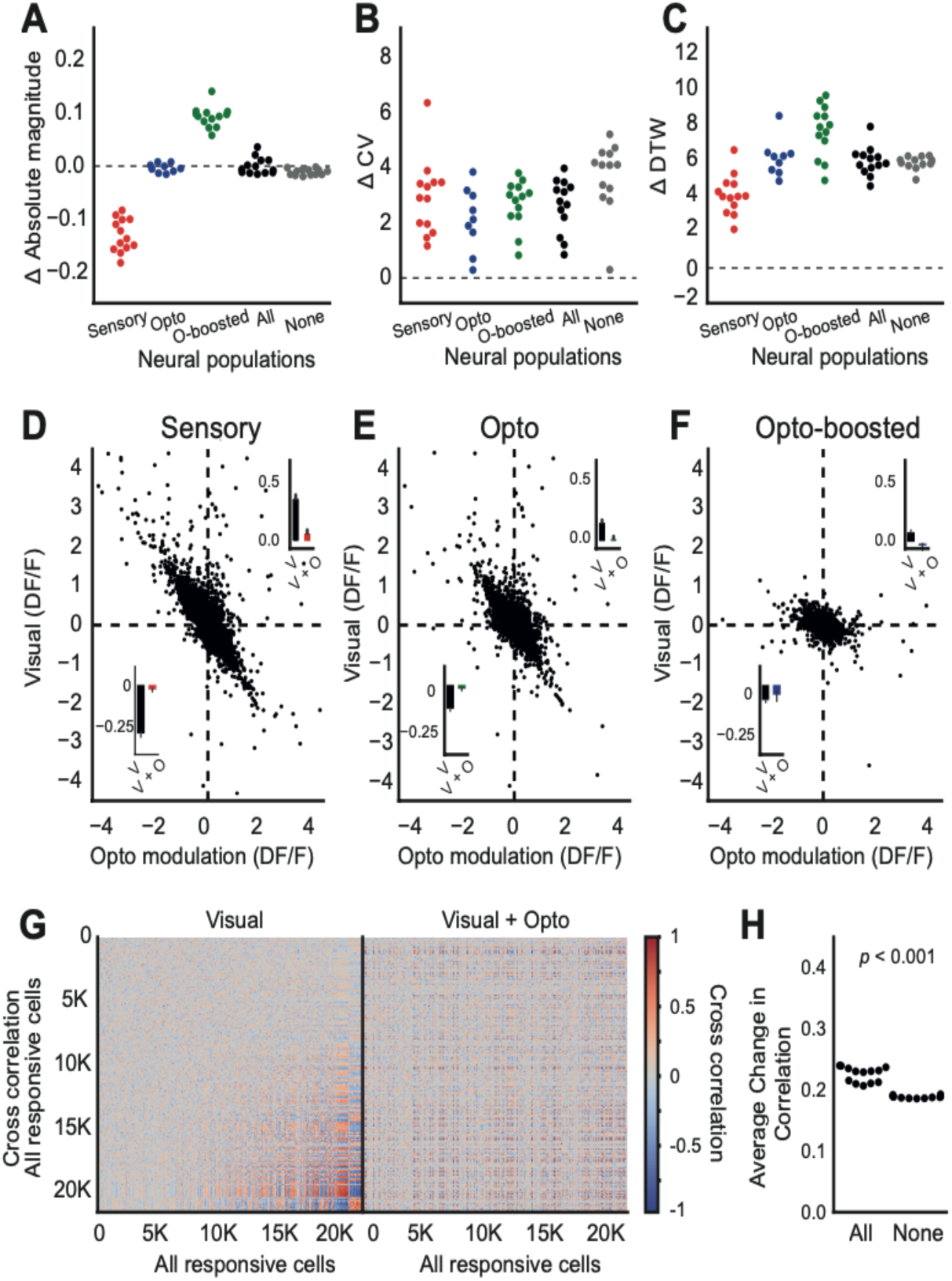
Modulation of claustrum axons enhances cross-correlation in the dPFC neural network. **A)** Swarm plot for the absolute magnitude of response changes for sensory, opto, opto-boosted, all (includes any responsive cells) and none (cells do not respond to any stimulation) neuronal populations. Each dot represents an individual animal (n = 13). **B)** Swarm plot for changes in the coefficient of variation (ΔCV) for each neuronal population. **C)** Swarm plot for dynamic time warping (ΔDTW) differences for each neuronal population. **D-F)** Scatter plots demonstrate the relationship between visual response (DF/F) and opto modulation (DF/F) in sensory (A), opto (B), and opto-boosted (C) neuronal subpopulations. The dotted lines indicate zero levels. **G)** Heatmaps display cross-correlation coefficients among all responsive cells during visual (left) and visual+opto (right) conditions. **H)** The average correlation changes after optostimulation, highlighting shifts in neural connectivity and synchronization under combined stimulation.

To assess the claustrum modulation on the dPFC neural network, we first examined the opto modulation compared to the neural response to visual cues. Correlation analysis revealed a strong negative correlation. As the ΔF/F response to visual stimuli increased, photostimulation had a more pronounced inhibitory effect. In contrast, when the ΔF/F response was negative, photostimulation led to an excitatory effect across all subpopulations (sensory: **Fig. 3D**, r = - 0.876, *p* < 0.001; opto: **Fig. 3E**, r = −0.851, *p* < 0.001; opto-boosted: **Fig. 3F**, r = −0.735, *p* < 0.001). Hence, claustrum modulation in dPFC cells strongly depends on the baseline state of responsiveness of the given cell, and this dependency is not limited to excitatory cells. Additional experiment revealed that inhibitory cells exhibit a similar pattern of modulation, indicating that this effect was consistent across different cell types (**Supplementary Fig. 3.1**, n = 5 Nkx2.1-GCaMP6s transgenic mice).

Then, we assessed the collective impact of all responsive cells including the three identified subpopulations by examining the cross-correlation of trial-averaged responses across all neurons. The cross-correlation matrix for visual stimuli showed distinct cell group clustering, suggesting a highly coordinated neural network (**Fig. 3G**, left). Conversely, the matrix for visual + opto stimuli demonstrated a more uniform distribution, with an absence of distinct clustering (**Fig. 3G**, right, using the same clustering as defined by the visual condition). When we calculated the average change in cross-correlation due to the optostimulation across animals, an increase in correlation was noted for all responsive as well as none responsive cells (**Fig. 3H**, significantly higher in all responsive cells than none responsive cells, *p* <0.001). Notably, while the coefficient of variation analysis indicated higher variability in responses across trials within individual cells, the cross-correlation analysis, which examines trial-averaged responses, revealed a more correlated response across the population. This indicates that the visual + opto stimuli facilitated a more homogenized and correlated interaction among all cells. This synergy among subpopulations likely contributes to refining dPFC activity, strengthening functional connections between neurons, and integrating them into a more cohesive network.

### Claustral modulation-induced network homogeneity is enhanced by experience

To investigate how claustral modulation affects neural flexibility driven by experience, we trained mice (n = 8) in a Pavlovian conditioning task with the same visual stimuli. Mice were water-restricted and presented with the same visual cues as in the experiments above followed by a drop of water (80% of the trials). 7 mice showed anticipatory licking activity before the reward within 10 days (**Fig. 4B-D**, *p* = 0.028), indicating that they had learned to associate the stimuli with the reward. After mice learned the association, they underwent the previous recording protocol in which they were passively exposed to visual cues with and without photostimulation. Notably, in this recording, the animals were familiar with visual stimuli although it was passively presented.

**Figure 4:**
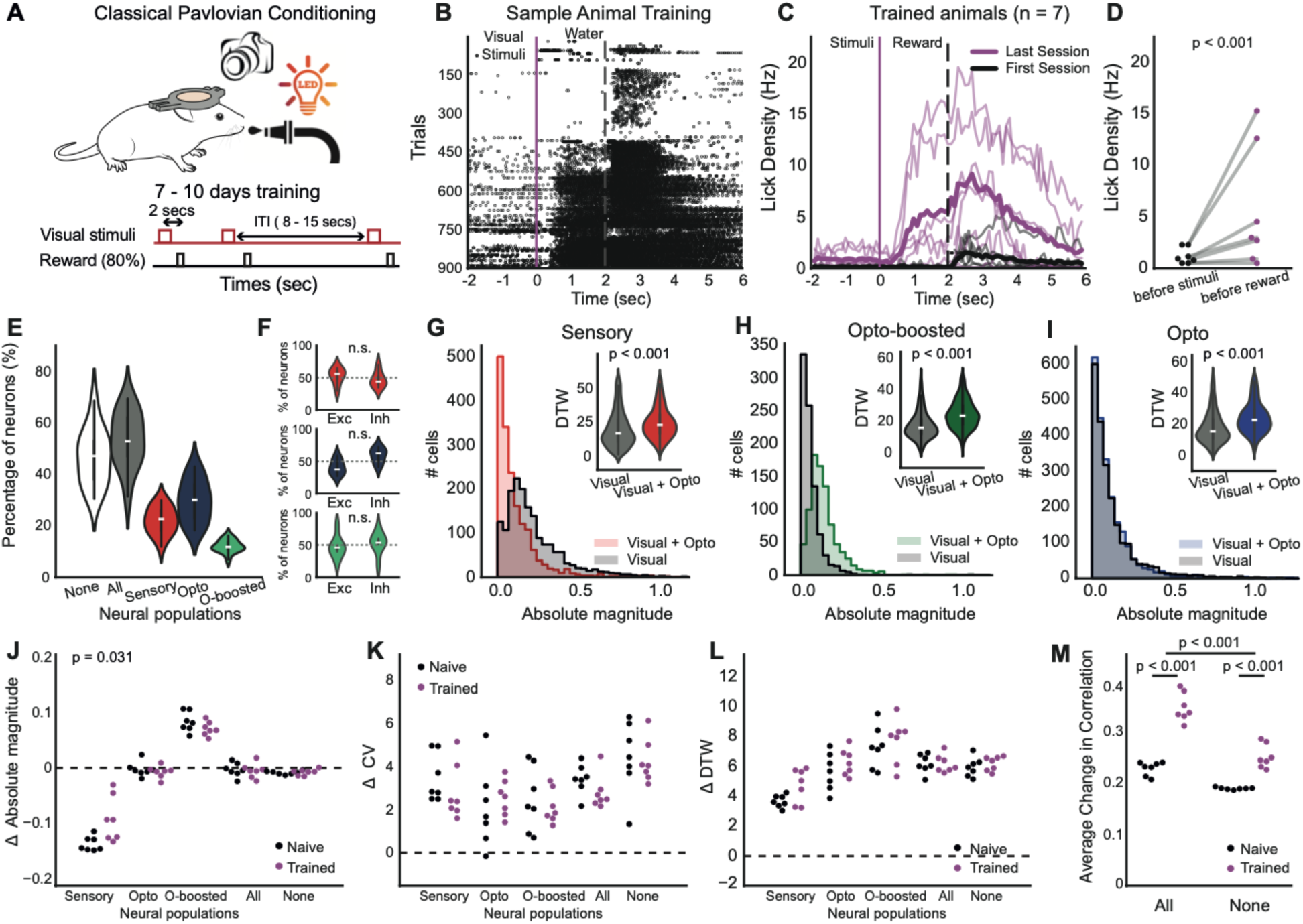
Claustral modulation-induced neural homogeneity is enhanced by training. **A)** Schematic illustrating the experimental design for classical Pavlovian conditioning in mice, involving visual stimuli over a 7-8 day training period with an 80% reward rate. **B)** A raster plot displaying lick responses from a sample animal during training, aligned to visual stimulus onset and reward delivery cues. **C)** Line plots presenting the average lick rate per second for trained animals (n = 7) across the first (black) and the last (magenta) session of the training. Thick line is for average, thin line is for individual mice. **D)** Average the lick density (1 sec) before stimuli and before reward. **E)** Violin plots depicting the percentage of responsive neurons within various neural populations: non-responsive (None), all responsive, sensory responsive, optogenetically responsive (Opto), and opto-boosted (O-boosted). **F)** Box plots comparing the percentage of excitation (Exc) and inhibition (Inh) responses among different neural populations.**G-I)** Absolute magnitude and dynamic time warping (DTW) analysis comparing sensory (G), opto-boosted (H), and opto (I) neural populations under visual only and visual+opto conditions. **J)** Swarm plot for the differences of absolute magnitude of neuronal responses to visual and visual+opto stim across neural populations in naive versus trained. Each dots represent one animal. **K)** Same with J but for coefficient of variation (ΔCV). **L)** Same with J but for ΔDTW. **M)** The average change in cross-correlation coefficients between response patterns of naive versus trained animals.

Similar to the naive animal recordings, we found three subpopulations in similar ratios for the trained animal recordings (53% responsive cells, 22% sensory responsive cells, 30% opto boosted cell and 12% opto-boosted cells, **Fig. 4E**). However, we observed no significant differences in excitatory and inhibitory responses among opto responsive cells, unlike the findings from the naive dataset (**Fig. 4F**, sensory: *p* = 0.578; opto *p* = 0.110; opto-boosted *p* = 0.297, pairwise t-test). These three subpopulations displayed similar patterns post-training;(i) sensory-responsive neurons exhibited reduced response magnitude with an increased disparity in neural response upon claustrum axon photostimulation (**Fig. 4G**), (ii) opto-boosted neurons showed amplified response magnitudes alongside greater neural variability (**Fig. 4H**), and (iii) optogenetically responsive neurons displayed consistent absolute magnitudes but demonstrated enhanced variance in neural responses (**Fig. 4I**).

The fundamental mechanisms across subpopulations remained consistent; the change in absolute magnitude (**Fig. 4J**, sensory *p* = 0.031; opto *p* = 0.110; opto-boosted *p* = 0.297, paired t-test), the change in the coefficient of variation ( **Fig. 4K**, sensory *p* = 0.219; opto *p* = 0.375; opto-boosted *p* = 0.578, paired t-test), and the change in DTW (**Fig. 4L**, sensory *p* = 0.0782; opto *p* = 0.469; opto-boosted *p* = 0.812, paired t-test) was greater than zero but did not differ in trained animals compared to naive animals across populations. This suggests that while training influences how visual cue perceived and behavioral responses, the intrinsic variability in neural responses induced by photostimulation remains relatively stable. Hence, the modulatory effects of the claustrum on these neurons are robust against experiential changes — at least for the aspects of neural response variability measured here.

On the other hand, correlation analysis revealed an average increase in correlation among neuronal responses in trained animals, indicating a more synchronized and integrated network response following training (**Fig. 4M**). A two-way ANOVA showed significant main effects of training (F (1,24) = 112.57, *p* < 0.001) and responsiveness (F(1,24) = 64.40, *p* < 0.001).

Additionally, there was a significant interaction between training and responsiveness (F (1,24) = 12.86, *p* = 0.002), indicating that the effects of training varied depending on cell responsiveness. This implies that experience amplifies the effects of claustrum axon stimulation, leading to more coordinated neural activity, particularly in responsive cells, while preserving the underlying mechanisms.

### Silencing the claustrum also enhances neural flexibility in the dPFC

To further elucidate the functional role of the claustrum, we conducted an additional photostimulation experiment to explore the effects of claustrum axon silencing on dPFC responses in both naive and trained animals. The same methodology was used as in the activation experiment, except we injected eOPN3 to silence axonal neural activity (**Fig. 5A-D**). Notably, eOPN3 is a light-sensitive opsin that functions as an inhibitory tool by reducing neuronal activity upon photostimulation. This contrasts with ChrimsonR, which we use for activation experiments, as ChrimsonR is a red-light-sensitive opsin that depolarizes neurons to induce excitation. Thus, while Chrimson activates neural circuits via depolarization, eOPN3 effectively silences them through hyperpolarization (activates a signaling cascade leading to hyperpolarization and subsequent silencing of neural axonal activity, providing an effective means of optogenetic inhibition (Mahn et al., 2021), see supplementary methods for more details)

**Figure 5:**
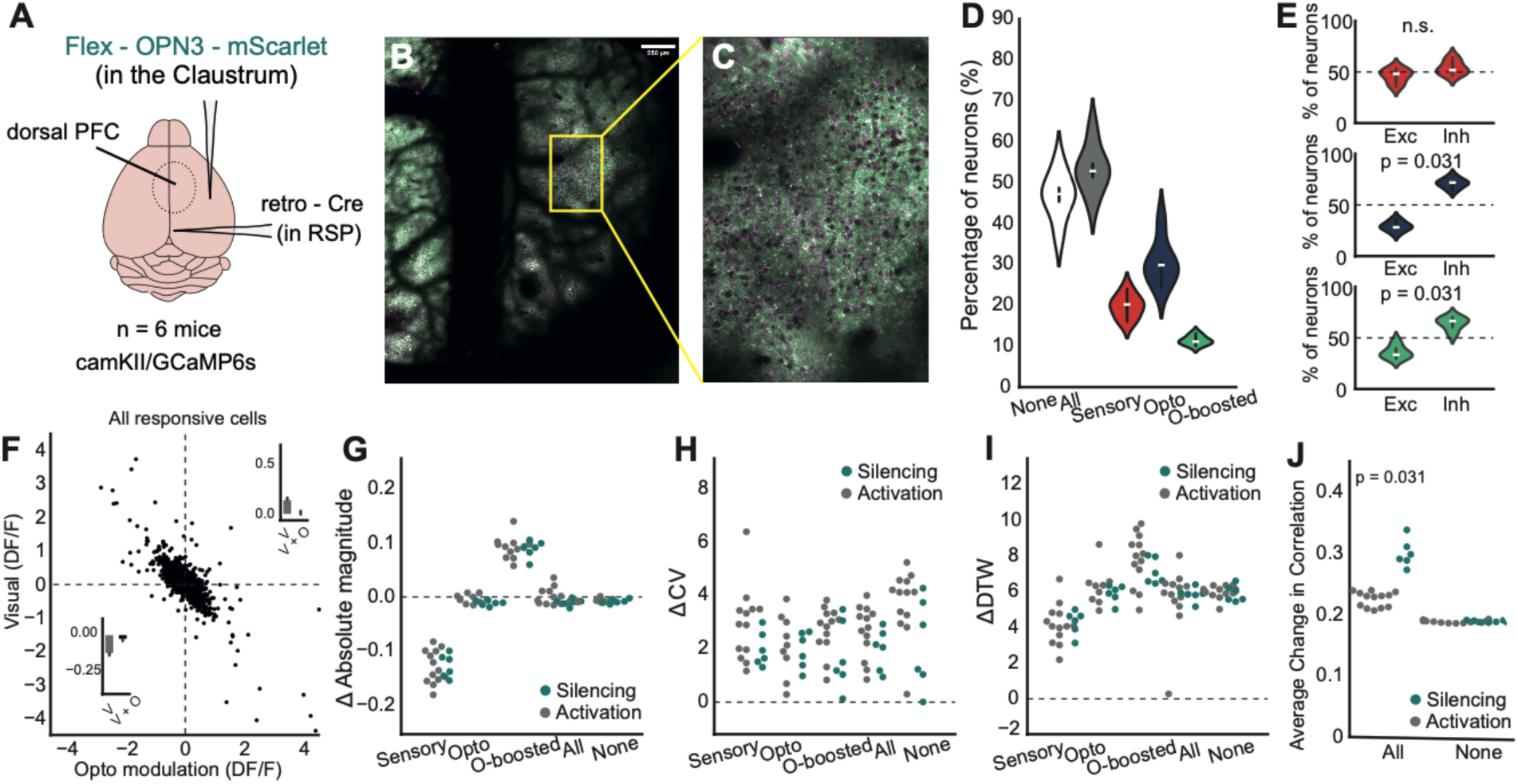
Claustrum axon silencing enhances network synchrony in the dPFC to a greater degree than activation. **A)** Schematic of the experimental setup illustrating the targeting of Flex-eOPN3-mScarlet expression in the claustrum and retro-Cre in the retrosplenial cortex (RSP) and cranial window over dPFC (n = 7 camKII/GCaMP6s mice) **B)** Representative confocal microscopy images showing axonal expression of Chrimson (red) in dPFC. **C)** Additional zoom confocal images showing the targeted axons within the dPFC. **D)** Violin plots displaying the percentage of responsive neurons across different neural populations: non-responsive (None), all responsive (All), sensory responsive, opto responsive, and opto-boosted neurons. **E)** Box plots comparing the percentages of excitatory (Exc) and inhibitory (Inh) responses within each neural population. **F)** Scatter plot for the relationship between visual response (dF/F) and opto modulation (dF/F) across all responsive neurons. **G)** Swarm plots for the absolute magnitude of neuronal responses across different groups, differentiated by ChrimsonR (activation) and eOPN3 (silencing) expression. **H)** Same with H but for coefficient of variation (ΔCV). **I)** Same with G but for dynamic time warping values (ΔDTW) **J)** The average change in cross-correlation coefficients between two groups.

Initially, we assessed the percentage of responsive neurons (**Fig. 5E**). The distribution of responsive neurons closely mirrored that observed during claustrum axon activation: 53% of the cells were responsive, with 20% classified as sensory-responsive, 30% as opto-responsive, and 11% as opto-boosted. Notably, in the opto-boosted subpopulation of naive animals (addition to opto responsive cells, *p* = 0.031), inhibited responses were more pronounced than excited responses, a difference not seen during claustrum axon activation (**Fig. 5F**, green plot for opto-boosted, *p* = 0.031). Moreover, the degree of the modulation effect was strongly correlated with the cell responsiveness (**Fig5F**, *r* = −0.999, *p* < 0.001)

Then, we explored each subpopulation dynamics and found similar dynamics to the activation experiment (**Fig. 5G**, absolute magnitude, sensory: *p* = 0.966; opto *p* = 0.066; opto-boosted *p* = 0.765; **Fig. 5H** ΔCV, sensory: *p* = 0.106; opto *p* = 0.776; opto-boosted *p* = 0.179; **Fig. 5I**, ΔDTW, sensory: *p* = 0.701; opto *p* = 0.456; opto-boosted *p* = 0.244). The only difference was a higher change in the cross-correlation for the silencing group than the activation group **Fig. 5K, A** two-way ANOVA, main effects of training: F (1,34) = 89.521, *p* < 0.001, responsiveness: (F(1,34) = 264.120, *p* < 0.001,interaction F(1,34) =87.529, *p* = 0.002), suggesting that silencing the claustrum enhances network synchrony in the dPFC to a greater degree than activation, though through similar mechanisms.

We then trained these mice on the same Pavlovian task and showed no differences between naive and trained data in terms of the measured neural dynamics (**Supplementary Fig. 5.1**). These findings suggest that the claustrum exerts a dual regulatory influence on the dPFC, modulating its output in a flexible manner depending on whether the claustrum is activated or silenced.

Overall, these findings suggest that the claustrum serves as a critical modulator of neural activity in the dPFC, influencing both the amplitude and variability of neuronal responses as well as the overall synchrony of the neural network. The effects observed during claustral silencing versus activation provide insights into the dual regulatory capacity of this brain region, highlighting its importance in enhancing neural flexibility to optimize brain functions.

## Discussion

Our results demonstrate the significant modulatory effect of claustrum axons on dPFC neurons during sensory processing. The diversity of responses observed across the sensory, opto-responsive, and opto-boosted subpopulations underscores the complex role the claustrum plays in shaping neuronal dynamics. Notably, the opto-boosted neurons, which exhibited no response to visual or optogenetic stimuli alone but responded to their combination, highlight the claustrum’s ability to selectively enhance neural responsiveness under certain conditions. This modulation, characterized by increased response variability and temporal dynamics, supports the hypothesis that the claustrum enables greater flexibility in sensory integration within the dPFC.

The differential response patterns between excitatory and inhibitory neurons further suggest that the claustrum’s influence is not uniform across the dPFC network. The finding that opto-responsive cells exhibited a stronger prevalence of inhibitory responses aligns with previous reports of the claustrum’s role in regulating inhibition within cortical networks (Jackson et al., 2018; Kim et al., 2016). The increased synchronization observed during the claustrum silencing experiment suggests that the loss of claustrum-mediated input leads to compensatory network adjustments, potentially to maintain functional coherence. This enhanced synchronization might reflect an intrinsic mechanism by which the dPFC preserves information processing capacity in the absence of normal claustral input.

Interestingly, training did not significantly alter the basic response properties of the identified subpopulations, suggesting that the claustrum’s modulatory effects are robust against experience-dependent plasticity in the context of sensory integration. However, the observed increase in network correlation post-training implies that while the claustrum supports flexibility and variability in neural responses, learning processes drive a more coordinated network state. This dual capability of the claustrum—to promote both variability and synchronization—suggests that it operates as a key regulator of both flexibility and stability in the neural circuitry of the dPFC, facilitating adaptive sensory processing.

The differential effects of claustrum activation and silencing in naive versus trained networks reveal significant insights into its functional role. While both activation and silencing of claustrum axons seem to produce similar outcomes in naive animals, activation uniquely enhances cross-correlation in trained networks, whereas inhibition does not alter the trained network’s dynamics. This suggests that the claustrum might exert an inhibitory influence under baseline conditions, as evidenced by the more pronounced inhibited responses in the opto-boosted subpopulation of naive animals, compared to excited responses—a distinction not observed during activation. These findings align with previous research, indicating that the claustrum may serve to diversify neural responses or avert excessive synchronization, which could otherwise lead to inflexible and less adaptable neural processing. When the claustrum’s modulatory effect is diminished, there appears to be a compensatory increase in synchronization, likely aimed at preserving coherent processing. Such adaptability in response dynamics highlights the claustrum’s pivotal role in enhancing neural flexibility, ensuring that the brain can swiftly adjust to varying conditions and demands.

There are two notable caveats in this approach. Firstly, the activation of claustrum axons in the PFC through widefield stimulation might not encompass all axons, as activation likely depends on the specific circuitry involved (Shelton et al., 2024; Erwin et al., 2021). Consequently, similar findings in both activation and silencing could diverge based on these selective activations. A potential study could involve activating the claustrum while recording from the PFC, although this remains challenging due to the claustrum’s elongated, sheet-like structure. Secondly, there are inherent differences in the response kinetics between eOPN3 and ChrimsonR, which are two distinct optogenetic tools. eOPN3 responds at a different rate and pattern compared to ChrimsonR, which could influence the interpretation of our results. Despite these differences, we chose to use the same photostimulation protocol for both to maintain consistency in the experimental setup. This approach ensures that any observed effects are due to the biological interactions rather than variations in stimulation techniques. However, it also means that the results should be viewed with an understanding of how these kinetic differences might impact the findings. By standardizing the protocol, we aimed to clearly isolate the contributions of the CLA-PFC interactions, recognizing that the kinetic variations might affect the response magnitude or timing. This highlights the importance of methodological consistency while also acknowledging the need to carefully consider the inherent properties of different optogenetic tools.

In conclusion, our findings underscore the claustrum’s role in dynamically modulating neural activity in the dPFC, enhancing the flexibility and synchronization of neuronal responses. The differential effects of activation and silencing of claustrum axons reveal its bidirectional regulatory capacity, emphasizing its importance in optimizing brain function. Future studies should explore the specific mechanisms through which the claustrum exerts this dual control and investigate its role in more complex cognitive tasks that require the integration of multiple sensory modalities.

## Materials & Methods

### Animals

Male and female C57BL/6J or Nkx2.1Cre;Ai9 background mice were used in these experiments. Twenty four GCaMP6s transgenic mice (CamKIIa-tTa x B6;DBA-Tg(tetO-GCaMP6s)2Niell/J) were used for imaging of the claustrum axon activation and inhibition experiments. Additionally, five Nkx2.1-Cre x tdTomato were used for inhibitory neuron imaging experiments. Mice were between 5-11 weeks of age when surgery was performed.Animal experimentation was carried out in accordance with the guidelines and regulations of the UK Home Office (Animals in Scientific Procedures Act of 1986) and the University of Oxford Animal Welfare and Ethical Review Board.

### Stereotaxic surgery

Mice were prepared for imaging experiments through a single surgical session that included headplate implantation, cranial window placement, and viral injection. Initially, the mice were anesthetized with 3% isoflurane, placed in a heated stereotaxic frame, and given intraperitoneal injections of 5 mg/kg meloxicam (Metacam) and 0.1 mg/kg buprenorphine (Vetergesic). They were maintained under 1.5% isoflurane and kept warm on a 37°C heating pad throughout the surgery. The scalp was sanitized with chlorhexidine gluconate and isopropyl alcohol (ChloraPrep), followed by the application of a local anesthetic (Bupivacaine) under the scalp. A midline incision was made in the scalp, which was then retracted to reveal the skull, leveled manually between the bregma and lambda landmarks. The site for the 4mm cranial window was marked stereotaxically (AP: −1 mm to 3 mm; ML: either −1 mm to 3 mm or −3 mm to 1 mm from Bregma, depending on the side of injection). A circular craniotomy was drilled at this marked location, and the skull piece was removed after saline application. The dura mater on the injected side was carefully excised with regular saline washing.

To specifically label claustrum axons in the PFC, AAV.hSyn.Cre.WPRE.hGH (Retro-cre, 0.70e+13 gc/mL, 90 nL, Addgene #105553) was injected into the retrosplenial cortex (AP: −3.0, ML:0.5, DV:-1.0) of all mice to label the retrosplenial cortex projecting claustrum cells. Then, we injected AAV5-Syn-FLEX-rc[ChrimsonR-tdTomato] (Flex-ChrimsonR, 1.20e+13 gc/mL, 250 nL, Addgene #62723) was injected into the claustrum (AP: 1.0, ML: 3.4, DV:-2.7) for activation experiments and pAAV-hSyn1-SIO-eOPN3-mScarlet-WPRE (FLEX-eOPN3, 1.20e+13 gc/mL, 250 nL, Addgene #125713) for axon silencing experiments. Injections were performed with a glass needle and automated nanoinjector (Nanoject II™ Drummond) at a rate of 100 nL/min. The needle remained in place for 10 minutes post-injection to promote viral diffusion. A cranial window, consisting of a 4 mm coverslip attached to a 5 mm coverslip, was then inserted into the craniotomy and sealed with cyanoacrylate (VetBond) and dental cement. An aluminium headplate with an imaging well centred on the window was then secured in place with dental cement (Super-Bond C&B, Sun-Medical). Post-surgery, mice were moved to a fresh cage, provided with meloxicam jelly for pain relief, and allowed a three-week recovery period to achieve optimal viral expression before beginning experiments.

### Awake head-fixed two-photon calcium imaging and widefield photostimulation

All two-photon imaging was performed using a two-photon microscope (Ultima 2pPlus, Bruker Corporation) controlled by Prairie View software (Bruker Corporation), and a femtosecond-pulsed, dispersion-corrected laser (Chameleon, Coherent). Total power was modulated with a Pockels cell (Conoptics) and maintained at or below 50 mW on a sample for all experiments. Imaging was performed using a Nikon 16X 0.8NA water immersion lens. The lens was insulated from external light using a custom 3D-printed cone connected to a flexible rubber sleeve. A wavelength of 920 nm and 50 mW power on the sample was used for visualizing GCaMP6s. An imaging rate of 30Hz and a 512×512 pixel square field of view (FOV) were used for all recordings.

Widefield (1P) optogenetics was performed in conjunction with 2P calcium imaging (as described above) using the widefield excitation module of the Bruker 2pPlus. Orange light photostimulation (595 nm, 10 x 25 ms pulses @ 40Hz, at 1.5mW power on a sample), was performed during 2P imaging. To avoid damage to the 2P imaging photomultiplier tubes (PMTs), high-speed shutters on all PMTs were triggered during light photostimulation. Triggers for light stimulation (Thorlabs M595L3 and LEDD1B LED driver) and the high-speed PMT shutters were provided to a using PackIO.

Once mice had completely recovered from surgery, animals were first acclimated to head fixation under the microscope. Next, ChrimsonR and eOPN3 expression levels in claustrum axons in PFC were assessed by eye. Animals in which no labelled axons could be found in the cranial window were excluded from future experiments. Animals with ChrimsonR or eOPN3 labelled axons in the cortex were then used for activation or inactivation experiments.

Sensory stimuli were delivered using a data acquisition card (National Instruments) and PackIO software. Briefly, custom MATLAB (MathWorks) code was used to generate voltage traces. These traces were then used by PackIO to output timed voltage from the data acquisition card to an LED (Thorlabs) for visual stimulation, that lasted 500ms.

During each experiment, mice were first head-fixed under the microscope. Imaging was performed in an enclosed hood to minimize visual stimuli, and white noise was used to obscure extraneous sounds. The surface of the cranial window was levelled relative to the imaging plane using a tip-tilt stage (Thorlabs). During each imaging session, mice were presented with 30 randomly interleaved stimulus presentations with and without optostimulation separated by randomly generated 8-15 second intertrial intervals (30 visual-only trials, 30 opto-only trials and 30 visual + opto trials). These 90 stimuli were randomly drawn. The order of each trial type was randomly generated each day. After all stimuli were delivered, a new FOV was then selected and the sensory stimulation and photostimulation were repeated.

### Behavioural training

Pavlovian conditioning was employed as the training task to familiarise the visual cue to the mice, utilizing a 2-second LED stimulus followed by a water reward in 80% of the trials. The LED was positioned unilaterally to ensure consistent exposure and response. Over a 7-10 day training period, animals were subjected to 150 trials per day, with each trial separated by an inter-trial interval (ITI) ranging from 8 to 15 seconds.

### Perfusion and tissue sectioning

The presence of claustrum labeling in each animal was confirmed using post-mortem coronal slices, spanning from 1.5 mm anterior to bregma to 0 mm, as shown in Supplementary Figure 1.1. Mice were deeply anesthetized with 5% isoflurane followed by an overdose of pentobarbital administered intraperitoneally. Subsequently, transcardial perfusion was performed using 0.01 M PBS, followed by fixation with 4% PFA. The brains were extracted and allowed to fix overnight in 4% PFA. After fixation, the brains were transferred to 0.01 M PBS and prepared for sectioning with a Leica VT1000S vibratome. Coronal slices, 100 μm in thickness, were obtained and either immediately mounted in 0.01 M PBS or stored in a tissue freezing solution (45% 0.01 M PBS, 30% ethylene glycol, 25% glycerol) at −20°C for up to three years. Before mounting, the tissue was washed three times for 5 minutes each in 0.01 M PBS and then coverslipped. Whole-slice and claustrum images were captured at 16X magnification using a 2P microscope equipped with a Coherent Vision-S laser, Bruker 2PPlus microscope, and a Nikon 16X 0.8 NA objective.

### Data Avaliablity

All processed data and the functions used to generate the figure panels in this study are available in GitHub (huriteatg/clapfcstimulation repo)

### Data Analysis

All analyses were performed with custom routines using Python 3.7.9 and open source packages unless otherwise stated.

#### 2-photon calcium imaging analysis

Calcium imaging data were preprocessed using Suite2P (Pachitariu et al., 2016) to remove motion artifacts and cell segmentation.We computed ΔF/F for each cell using the equation:

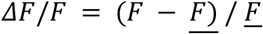

where <u>*F*</u> represents the mean of *F* across time through the entire session. For cell ROIs selected by suite2P, *F* was first corrected for neuropil fluorescence by subtracting 0.7*FNeu. After calcium traces were exported from Suite2P, all analyses were carried out using custom Python and Matlab code.

Extracted calcium signals were then analyzed to identify cells that significantly responded to visual, opto or visual + opto stimulation. Significantly responsive cells were identified by using a non-parametric Mann-Whitney U test to compare the signal in the 1 second before and after stimulus onset. Multiple comparisons correction was performed using the Benjamini-Hochberg false discovery rate analysis with an alpha of 1%.

#### Event time extractions

Synchronised timeseries data signals collected as individual channels in PackIO (visual cues and photostimulation) were processed with custom code. Widefield photostimulation timings were retrieved from the high-speed shutter loopback signal collected as a temporally synchronised PackIO channel. The photostimulation trigger timestamp and the total duration of the photostimulation protocol were used to define each photostimulation trial’s onset and offset. The photostimulation onset/offset timestamps were cross-referenced to the imaging frame clock signal to define photostimulation onset and offset in the imaging data. We excluded imaging frames between onset and offset of each photostimulation trial as photostimulation laser induced imaging artefacts and aberrant neuronal activity.

To quantify the intricacies of neuronal activity, we used four different metrics, absolute magnitude, coefficient of variation (CV), dynamic time warping (DTW), and empirical cumulative distribution function (ECDF). The absolute magnitude is calculated to determine the intensity of response changes by comparing mean fluorescence levels before and after a stimulus. This measure, by accounting for noise through standard deviation, aims to quantify signal changes from background fluctuations. The CV, which quantifies the variability of the fluorescence signal relative to its mean for each trial, providing insights into the consistency of neuronal responses across trials. The dynamic time warping (DTW) is employed to assess temporal dynamics by calculating the average distance between fluorescence response patterns across trials for each cell. This method captures the temporal similarities and differences, offering a deeper understanding of response timing and synchronization. Finally, The ECDF further by illustrating the cumulative distribution of fluorescence data, allows us to visualize and interpret the variability and distribution of responses across different conditions. Together, these calculations enable a comprehensive analysis of the data, highlighting both the magnitude and variability of neuronal responses as well as their temporal coordination. Together, these calculations enable a comprehensive analysis of the data, highlighting both the magnitude and variability of neuronal responses as well as their temporal coordination and distribution characteristics.

## Supplementary Figures

**Supplementary Figure 1.1:**
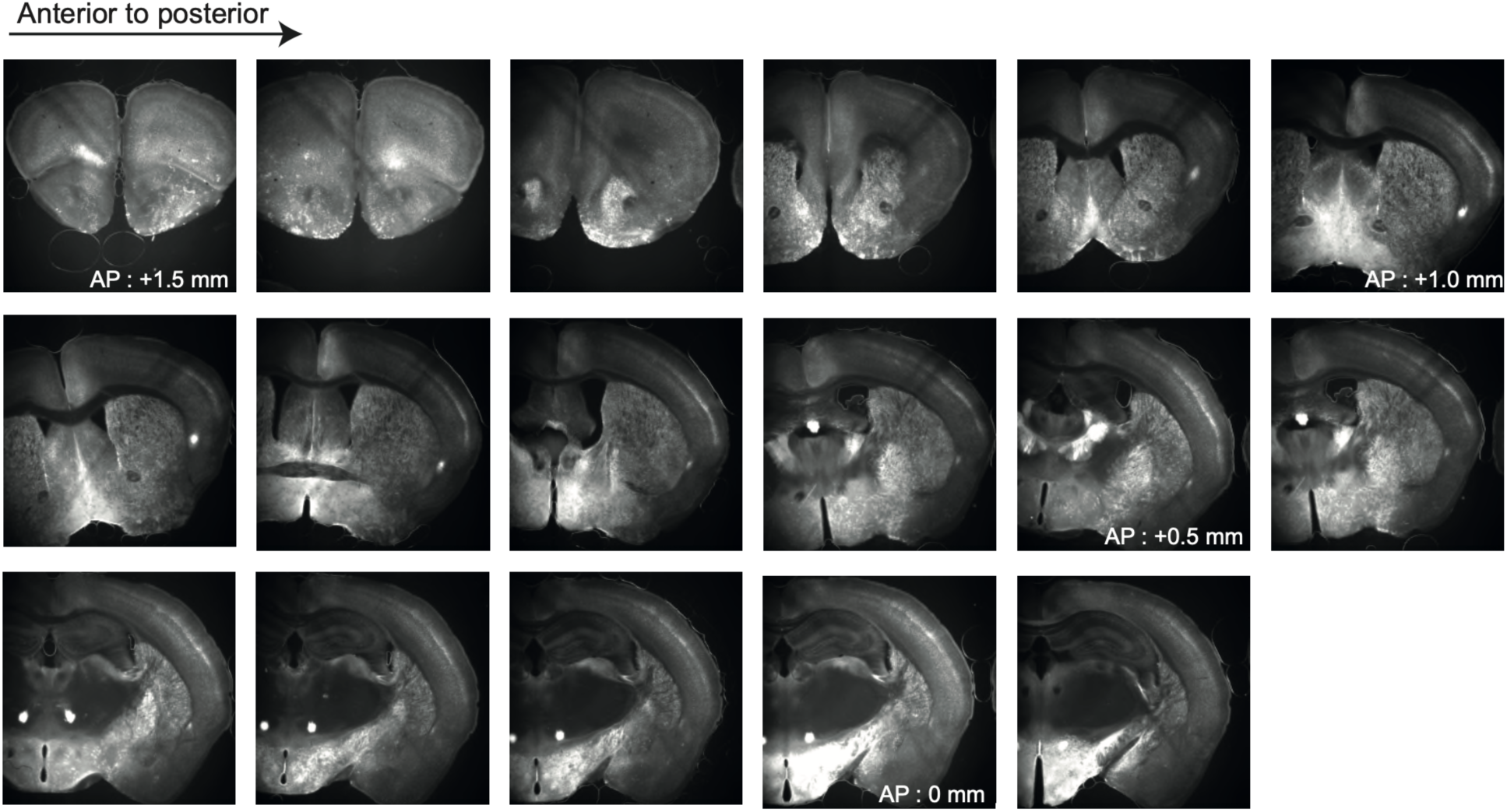
Coronal brain sections illustrating the distribution of claustrum cell labeling across anterior-posterior sections of the claustrum. The images are organized sequentially from anterior to posterior, with the first image on the left representing the most anterior section and the last image on the right representing the most posterior section. The sections start at approximately 1.5 mm anterior to the bregma, and claustrum labeling is evident from 1.2 mm, if not before, to 0.2 mm anterior to the bregma, covering nearly 1mm of the claustrum.

**Supplementary Figure 1.2:**
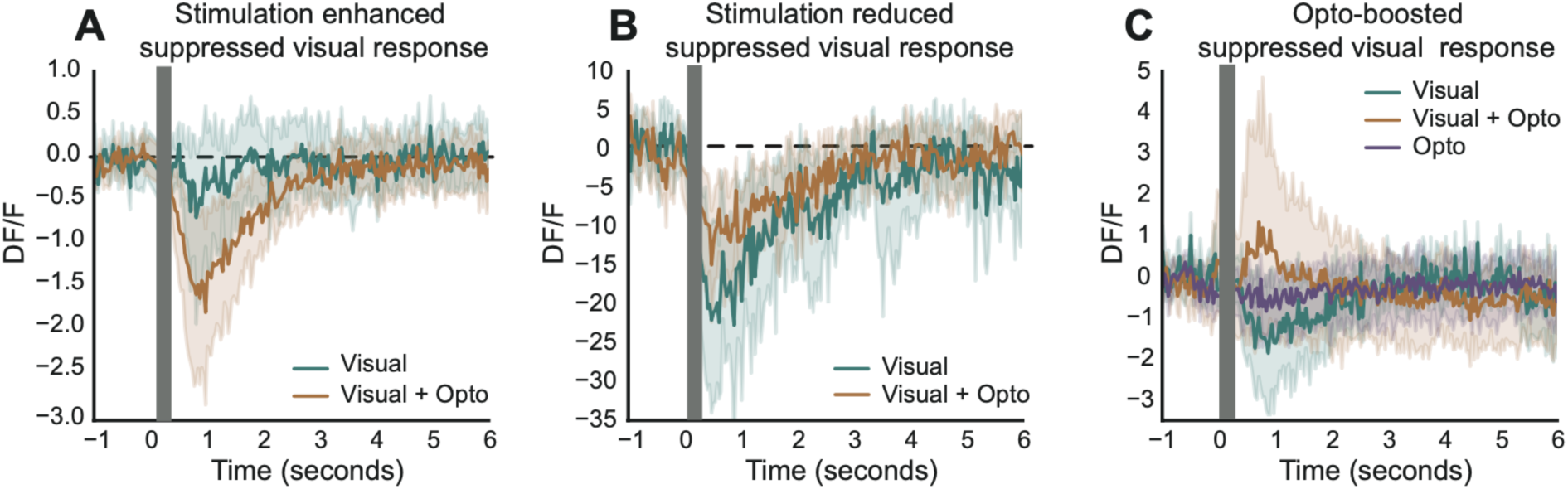
Various neural modulation with the claustrum axon activation through photo-stimulation. The time course of neuronal calcium responses (ΔF/F) illustrates the modulatory effects of the claustrum axon photostimulation on dPFC neuron responses to visual stimuli. Individual plots show different modulatory effects on suppressed visual responses when combined with optogenetic stimulation. Animals with ChrimsonR

**Supplementary Figure 2.1:**
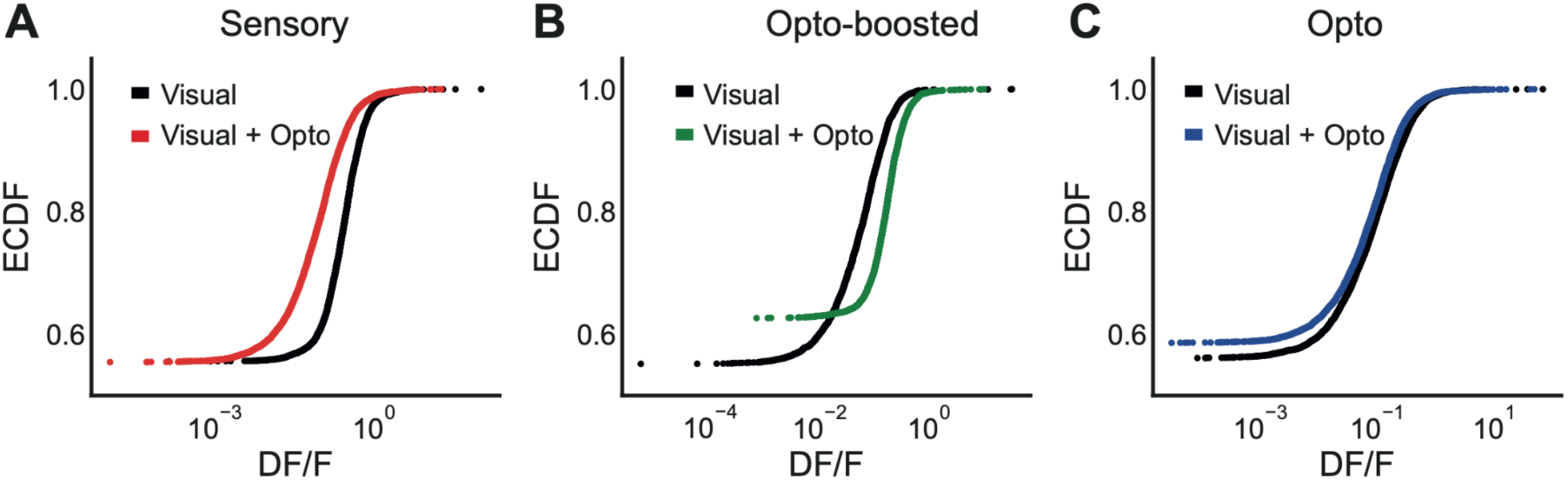
Empirical cumulative distribution function (ECDF) analysis of neural responses to combined visual stimuli presentation and axon activation through photostimulation. A) The ECDF plot shows the neural responses under different conditions. The visual + optogenetic stimulation (Visual + Opto) curve (red) is consistently above the visual-only curve (black) across all points, indicating broader neuronal engagement when visual stimuli are combined with optogenetic activation. B) The ECDF curves for opto-boosted neurons reveal a more rapid increase in the visual + optogenetic condition, suggesting that more neurons quickly reached the response threshold with optogenetic activation. The steeper rise of the Visual + Opto curve (green) implies that neurons achieve the specified response level more swiftly, highlighting enhanced neural activity at the collective level for opto-boosted neurons. C) The ECDF plot for the optogenetic-only condition (blue) shows the response in the presence of optogenetic stimulation alone compared to visual-only stimulation (black).

**Supplementary Figure 3.1:**
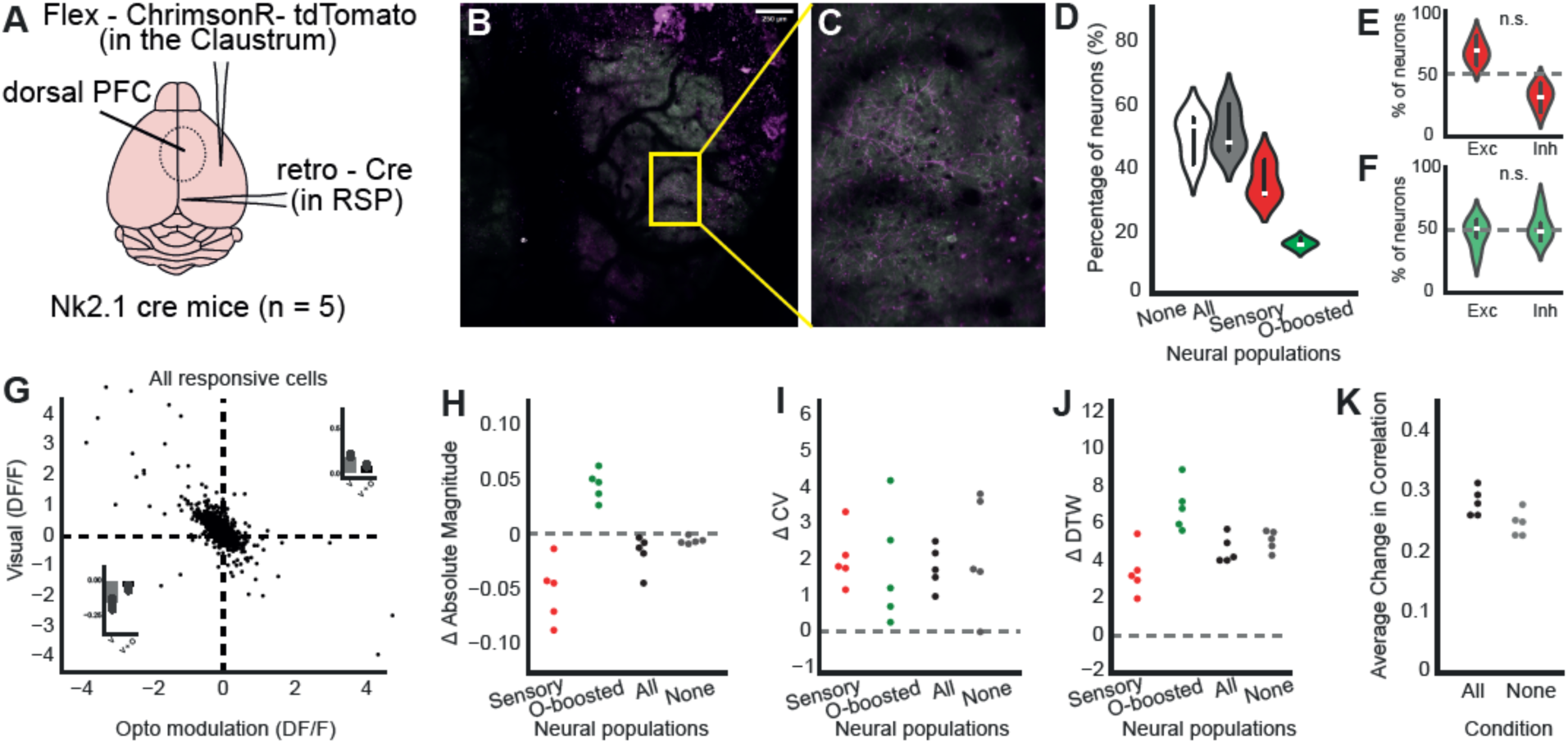
Claustrum axon activation enhances neural flexibility in the inhibitory cells and enhance homogeneity in dorsal prefrontal cortex. **A)** Schematic of the experimental setup for two-photon imaging of dorsal inhibitory PFC neurons expressing GCaMP6s (n = 5) in Nk2.1 mice. **B)** Stitched two-photon image of the dorsal surface of the brain showing GCaMP-expressing neurons (green) and claustrum axons labelled with tdTomato (magenta) within the imaging field of view. In this experiment, we presented visual and visual+opto stimuli, but did not include opto-only stimuli, so that there was no opto responsive subpopulation. **C)** The yellow box in panel B highlights the region magnified in panel C, where individual neurons are clearly visible. **(D)** Violin plot showing the percentage of neurons categorized into different neural populations. **(E-F)** Violin plots comparing the percentage of excitatory (Exc) and inhibitory (Inh) neurons within the neural populations categorized as Sensory and Opto-boosted, respectively. The n.s. (not significant) label indicates no significant difference between the proportions of excitatory and inhibitory neurons. **(G)** Scatter plot illustrating the relationship between visual response (ΔF/F) and opto modulation (ΔF/F) for all responsive cells. The plot shows a strong negative correlation between visual and optogenetic modulation (*r* = −0.521, *p* < 0.001) **(H)** Dot plot showing the change in absolute magnitude (Δ Absolute Magnitude) of neural responses across different neural populations under visual+opto stimuli. Sensory (*p* = 0.019 and opto-boosted (*p* < 0.001, paired t-test) subpopulation showed significant modulation in different directions. **(I)** Dot plot showing the change in the coefficient of variation (Δ CV) across the same neural populations, indicating variability in response magnitude across trials (sensory: *p* = 0.792; opto-boosted: *p* = 0.712, paired t-test). **(J)** Dot plot showing the change in Dynamic Time Warping (Δ DTW), a measure of temporal variability, across the same neural populations (sensory: *p* = 0.020; opto-boosted: *p* = 0.058, paired t-test). **(K)** Dot plot showing the average change in correlation across all conditions (All, None), highlighting the overall effect of optogenetic modulation on neural network coordination in the dPFC. Overall, the responses of inhibitory cells were modulated with claustrum axon modulation in a manner similar to that of CamKII-expressing cells

**Supplementary Figure 5.1:**
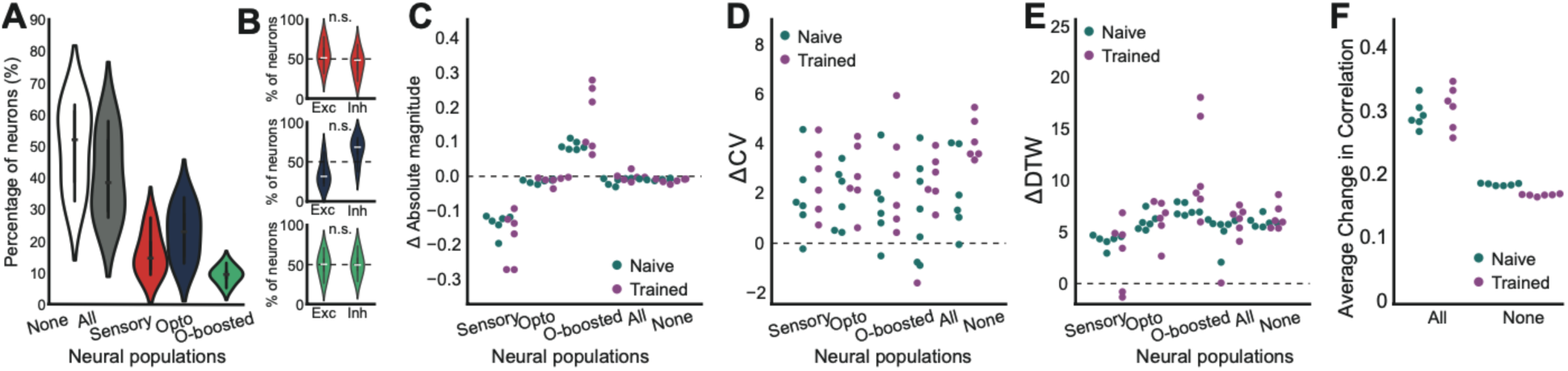
Training did not change the claustrum axon silencing-induced neural flexibility and network homogeneity. A) Violin plots displaying the percentage of responsive neurons across different neural populations: non-responsive (None), all responsive (All), sensory responsive, opto responsive, and opto-boosted neurons. B) Box plots comparing the percentages of excitatory (Exc) and inhibitory (Inh) responses within each neural population. C) Swarm plots for the absolute magnitude of neuronal responses across before (Naive) and after training (Trained). D) Same with C but for coefficient of variation (ΔCV). E) Same with C but for dynamic time warping values (ΔDTW). E) The average change in cross-correlation coefficients fo naive and trained data. The fundamental mechanisms across subpopulations remained consistent after the training; the change in absolute magnitude (**Fig. 5.1C**, sensory *p* = 0.219; opto *p* = 0.438; opto-boosted *p* = 0.156, paired t-test), the change in the coefficient of variation (**Fig. 5.1D**, sensory *p* = 0.563; opto *p* = 0.438; opto-boosted *p* = 0.562, paired t-test), and the change in DTW (**Fig. 5.1E**, sensory *p* = 0.563; opto *p* = 0.844; opto-boosted *p* = 0.063, paired t-test) was greater than zero but did not differ in trained animals compared to naive animals across populations. The correlation analysis revealed no change in correlation among neuronal responses in trained animals compared to naive animals (**Fig. 5.1F**). A two-way ANOVA showed no significant main effects of training (F (1,20) = 0.112, *p* = 0.741 (responsiveness: F (1,20) = 222.034, *p* < 0.001); interaction F (1,20) = 2.753, *p* = 0.113).

